# Bioresorbable Mesenchymal Stem Cell–Loaded Electrospun Polymeric Scaffold Inhibits Neointimal Hyperplasia Following Arteriovenous Fistula Formation in a Rat Model of Chronic Kidney Disease

**DOI:** 10.1101/2022.11.21.517369

**Authors:** Allan John R. Barcena, Joy Vanessa D. Perez, Marvin R. Bernardino, Jossana A. Damasco, Andrea Cortes, Huckie C. Del Mundo, Erin Marie D. San Valentin, Carleigh Klusman, Gino Martin Canlas, Francisco M. Heralde, Rony Avritscher, Natalie Fowlkes, Richard R. Bouchard, Jizhong Cheng, Steven Y. Huang, Marites P. Melancon

**Affiliations:** Department of Interventional Radiology, The University of Texas MD Anderson Cancer Center, Houston, TX 77030, USA; College of Medicine, University of the Philippines Manila, Manila 1000 Philippines; Baylor College of Medicine, Houston, TX 77030, USA; Department of Chemistry, Lamar University, 10009 Beaumont, TX 77710; Department of Veterinary Medicine and Surgery, The University of Texas MD Anderson Cancer Center, Houston, TX 77030, USA; Department of Imaging Physics, The University of Texas MD Anderson Cancer Center, Houston, TX 77030, USA; Section of Nephrology, Department of Medicine, Selzman Institute for Kidney Health, Baylor College of Medicine, Houston, TX 77030, USA; The University of Texas MD Anderson Cancer Center UTHealth Houston Graduate School of Biomedical Sciences, Houston, TX 77030, USA

**Keywords:** arteriovenous fistula, end-stage renal disease, mesenchymal stem cell, polymer, positron emission tomography, ultrasonography

## Abstract

In the setting of chronic kidney disease (CKD), the periadventitial injection of mesenchymal stem cells (MSCs) has shown significant potential in improving arteriovenous fistula (AVF) maturation by inhibiting neointimal hyperplasia (NIH). However, the rapid clearance of MSCs remains a challenge. Hence, we fabricated an electrospun perivascular scaffold from polycaprolactone (PCL) to support MSC attachment and allow gradual MSC elution at the outflow vein, the AVF site most prone to NIH. We performed a 5/6^th^ nephrectomy to induce CKD in Sprague-Dawley rats, followed by direct AVF formation and perivascular scaffold application. We then compared the following groups of CKD rats: no perivascular scaffold (i.e., control), PCL scaffold, and PCL+MSC scaffold. On ultrasonography, the PCL and PCL+MSC groups showed significantly reduced wall thickness and wall-to-lumen ratio and increased luminal diameter and flow rate. Of note, the PCL+MSC group showed greater improvement in luminal diameter and flow rate compared to PCL alone. Moreover, ^18^F-fluorodeoxyglucose positron emission tomography showed that only PCL+MSC resulted in a significant reduction in uptake. On histology, the PCL and PCL+MSC groups showed significantly reduced neointima-to-lumen and neointima-to-media ratios and reduced neointimal CD45, α-SMA, and vimentin fluorescence staining compared to the control. Although the two scaffold treatments did not differ significantly in histology, in vivo imaging suggested that the addition of MSCs promoted greater luminal expansion and blood flow and reduced the inflammatory process underlying NIH. Our results demonstrate the utility of mechanical support loaded with MSCs at the outflow vein immediately after AVF formation to support maturation by minimizing NIH.

## INTRODUCTION

Approximately 5.7 million individuals worldwide suffer from end-stage renal disease.^1^ In the United States, individuals with this condition have reached over 800,000.^2^ Globally, hemodialysis is still the most common method for replacing kidney function.^1^ For vascular access for hemodialysis, an arteriovenous fistula (AVF) is preferred over an intravenous catheter or an arteriovenous graft because of the lower risk of infection and fewer interventions to maintain patency.^3^ However, failure of an AVF to mature for hemodialysis use is reported to occur in up to 60% of patients.^4–8^ Salvage interventions for AVF stenosis, such as balloon angioplasty and stent placement, are available, but these interventions add to the treatment cost and patient morbidity.^8,9^ Furthermore, patients with AVFs requiring these interventions have decreased cumulative overall survival.^10^

One of the major pathologic etiologies associated with AVF maturation failure is neointimal hyperplasia (NIH), a cellular process that results from the synergistic action of inflammation, hypoxia, and hemodynamic stress on the anastomotic tissue. These pathways together promote endothelial cell proliferation and inward migration of pro-inflammatory cells, smooth muscle cells, and myofibroblasts, which result in vascular stenosis and thrombosis.^11,12^ To suppress NIH, several systemic therapies have been studied, including aspirin, clopidogrel, dipyridamole, warfarin, sulfinpyrazone, HMG-CoA reductase inhibitors, and fish oil. However, no systemic therapy has shown a clear benefit in improving AVF maturation rates,^13^ and a major limitation of systemic agents is dose-limiting toxicity. Localized interventions have the potential to improve AVF maturation rates by providing a direct therapeutic effect at the anastomotic site without the risk of systemic toxicity.^14^

The use of locally delivered mesenchymal stem cells (MSCs), which have immunomodulatory properties that help restore homeostasis, repair damage, and regulate cellular proliferation following vascular injury,^15,16^ has recently been explored. Yang et al.^17^ investigated the periadventitial injection of human adipose tissue–derived MSCs into the outflow vein of the AVF in immunodeficient B6.Cg-Foxn1nu/J mice and found that this approach significantly reduced the expression of known NIH mediators, such as monocyte chemotactic protein 1 (62%, *P* = 0.029) and hypoxia-inducible factor 1 alpha (62%, *P* = 0.0005) after 21 days. This strategy also significantly affected the outflow vein via increased mean luminal area (415%, *P* = 0.011), reduced mean neointimal area (77%, *P* = 0.013), and reduced neointimal cell density (83%, *P* < 0.0001).^17^ However, the washout of cells from the delivery site remains a challenge, as AVF maturation in humans can require 6-12 weeks.

One strategy to improve MSC retention and survival is the use of a perivascular scaffold that would provide mechanical support to the maturing AVF while also serving as an organic depot for MSCs to release regulatory substances to suppress NIH over time. The utility of an extravascular, non-resorbable scaffold (VasQ™, Laminate Medical Technologies, Israel) to improve AVF maturation and patency has been shown in a small single-center patient cohort.^18^ This solution, however, is only mechanical in nature. Polymeric scaffolds allow drugs and cell-based therapies to be delivered locally,^19^ providing an additional dimension to this prophylactic treatment schema. Several prior studies have shown that bioresorbable perivascular scaffolds loaded with antiproliferative agents can decrease NIH and improve AVF patency,^20,21^, but this strategy has not yet been utilized to deliver MSCs.

Through the process of electrospinning, bioresorbable synthetic polymers can be used to produce fibrous scaffolds that can mimic the three-dimensional architecture of collagen fibrils in the vascular extracellular matrix. Polycaprolactone (PCL) is a highly tunable and relatively affordable FDA-approved synthetic biodegradable polymer used in a wide variety of tissue-engineering applications due to its biocompatibility. Electrospun PCL has been used to generate engineered cardiac, tracheal, skeletal, integumentary, and vascular tissues.^22^ We have previously shown that an electrospun PCL scaffold can support the attachment and proliferation of MSCs.^23^ The objective of the current study was to determine the effects on NIH of PCL scaffolds with and without MSC attachment in a rat chronic kidney disease (CKD) model.

Ultrasonography (US) is used clinically to assess parameters that predict AVF maturation, such as luminal diameter and flow rate.^24^ One major disadvantage of US, however, is that the anatomic and physiologic variables may not correlate with underlying pathophysiology. ^18^F-fluorodeoxyglucose positron emission tomography (^18^F-FDG-PET) is a readily available imaging modality used in clinical practice to evaluate for sources of increased ^18^F-FDG uptake, such as inflammation, infection, and malignancy. In ^18^F-FDG PET, inflammation is detected when infiltrating inflammatory cells avidly take up the injected radiolabeled glucose, making it a widely used modality in identifying and monitoring inflammatory disorders. A recent case report showed that ^18^F-FDG-PET was successfully used to identify high ^18^F-FDG uptake at a patient’s suspected AVF infection.^25^ However, to the best of our knowledge, the utility of ^18^F-FDG-PET for measuring inflammation at the AVF anastomosis has not been assessed. Thus, a second objective of the current study was to validate the role of ^18^F-FDG PET in AVF inflammation. Overall, our study aims to provide new strategies for AVF patency support and assessment.

## METHODS

### Scaffold synthesis via electrospinning

The perivascular scaffolds were fabricated from PCL (average M_n_ 80,000, Sigma-Aldrich, St. Louis, MO) using a Spraybase electrospinning system (Avectas, Maynooth, Ireland). The polymer was dissolved in chloroform/methanol (9:1), and each sample solution was pumped through the syringe at a constant rate of 1.0 mL/h to the 20-gauge blunt-end needle tip. The distance between the needle tip and the mandrel collector was 15 cm. The polymer concentration and electrospinning voltage were optimized to produce aligned fibers that would facilitate cell-scaffold interaction and directional cell proliferation.

### Scaffold characterization

The fiber alignment, morphology, diameter, and pore size of each sample were determined using a Nova NanoSEM scanning electron microscope (Field Electron and Ion Company, Hillsboro, OR) with an EDAX energy dispersive spectroscopy (EDS) system (Ametek, Berwyn, PA). At least 100 segments of fibers and pore areas were randomly chosen and measured per sample using ImageJ (National Institutes of Health, Bethesda, MD). The porosity of the scaffolds was determined using the calculation method^26^ and the liquid intrusion method.^27^ The melting temperature (T_m_) and glass transition temperature (T_g_) of the electrospun scaffolds were determined using an STA PT 1000 thermogravimetric analyzer (Linseis, Selb, Germany). The ultimate tensile strength of the tubular scaffolds was determined using an MTESTQuattro materials testing system (ADMET, Norwood, MA) at ambient conditions.

### Cell culture and MSC attachment

Bone-marrow-derived Sprague-Dawley rat mesenchymal stem cells that express red fluorescent protein (RFP-MSCs) (Catalog no. CSC-C1315, Creative Bioarray, Shirley, NY) were cultured using DMEM with 10% FBS and 1% penicillin-streptomycin (Corning, Corning, NY). The scaffolds were subjected to ethylene oxide gas sterilization and then seeded with 1 × 10^5^ RFP-MSCs at 3^rd^ passage. After 12 hours, the samples were washed with PBS to remove unattached cells. For in vitro visualization of attachment, the scaffolds were imaged using an Eclipse Ti2 fluorescence microscope (Nikon, Melville, NY). For in vivo efficacy studies, cells were kept in culture media for a maximum of 24 hours prior to surgical application.

### AVF creation and application of the perivascular wrap

All rat experiments were approved by the Institutional Animal Care and Use Committee. We created a CKD model using Sprague-Dawley rats based on the two-step 5/6^th^ nephrectomy procedure by Wang et al.^28^ The first surgery involved the removal of both poles of the left kidney. The second surgery, performed 7 days after the first surgery to ensure rat survival, involved the removal of the entire right kidney. Creatinine and blood urea nitrogen (BUN) were obtained after the second procedure to confirm the decline in kidney function. We then created an AVF model via surgical anastomosis of the external jugular vein with the common carotid artery in an end-to-side fashion based on a modified procedure by Wong et al.^29^ AVFs were created in non-CKD rats and CKD rats 4 weeks after subtotal nephrectomy. In subsequent experiments, CKD rats were divided into three groups—control (i.e., no scaffold), PCL alone, and PCL+MSC. In the scaffold groups, following AVF formation and hemostasis, the sterilized polymer scaffold (1 mm x 5 mm) was wrapped around the outflow vein. The PCL+MSC group received a scaffold seeded with 1 × 10^5^ MSCs.

### Ultrasonography

The successful creation of an AVF was confirmed by the presence of pulsatile arterial waveforms in the external jugular vein visualized using the Vevo 2100-LAZR system (VisualSonics, Toronto, ON, Canada). Four weeks after surgery, the rats were anesthetized and imaged. Vevo B-mode, color Doppler, and pulse wave tool were used to assess the wall thickness, luminal diameter, and flow rate of the AVFs. A 15-MHz B-mode probe was used to get 6 measurements of the wall thickness and luminal diameter across the outflow vein. Wall-to-lumen ratio was calculated by dividing the average wall thickness by the average luminal diameter. An 18-MHz Doppler probe was used to measure flow velocity. The flow rate was derived from the flow velocity and average luminal diameter.

### ^18^F-fluorodeoxyglucose positron emission tomography

^18^F-FDG-PET/computed tomography (CT) images were acquired in Sprague-Dawley rats at baseline (i.e., non-CKD) and 4 weeks following the two-stage surgery (i.e., CKD). ^18^F-FDG PET/CT images were obtained using an m.CAM gamma camera (Siemens, Malvern, PA). The radioactive tracer—37 MBq of ^18^F-FDG in 200 μL of saline—was injected intravenously. Forty minutes after injection of the radioisotope, a 20-minute PET/CT scan was acquired of the AVF for each rat. The scans were reconstructed by a 2D maximum-likelihood expectation-maximization algorithm. Default reconstruction parameters were applied. The maximum standard uptake values (SUV_max_) of the AVFs were quantified.

### Histological analysis

Rats were euthanized by CO_2_ asphyxiation and thoracotomy after imaging. The AVFs were harvested after perfusion for epifluorescence visualization and paraffin embedding. To monitor the retention of RFP-MSCs at 2 and 4 weeks against perivascular injection of MSCs, the harvested vessels were fixed in 10% formaldehyde for 15 minutes, stained with 4’,6-diamidino-2-phenylindole (DAPI), and viewed using an Eclipse Ti2 fluorescence microscope (Nikon, Melville, NY). After the samples were embedded and sliced, the sections were divided for hematoxylin and eosin (H&E) and immunofluorescence multiplex staining with the following markers: (1) CD45, which is highly expressed in leukocytes and a marker of inflammatory infiltration; (2) alpha smooth muscle actin (α-SMA), highly expressed in smooth muscle cells but also expressed in fibroblasts; (3) vimentin, highly expressed in fibroblasts but also expressed in vascular smooth muscle cells and lymphocytes; and (4) DAPI. An Aperio LV1 real-time digital pathology system (Leica Biosystems, Buffalo Grove, IL) was used to capture images from the H&E slides, and quantification of the lumen, neointima, and media was done using ImageJ software (NIH). A Leica Versa fluorescent system (Leica Biosystems) was used to scan and capture images from the immunofluorescence slides, while the Halo image analysis platform (Indica Labs, Albuquerque, NM) was used to perform fluorescence cell counting.

### Statistical analysis

Quantitative data were presented as means ± standard deviations and analyzed using oneway analysis of variance (ANOVA) or a two-tailed *t* test. Statistical significance was defined as *P* < 0.05.

## RESULTS

### Physicochemical properties of electrospun PCL

Polymer concentration and electrospinning voltage were optimized to produce scaffolds with aligned fibers (Supplementary Figure S1). Figure 1 shows the representative morphology, diameter, and pore size of the selected PCL scaffold, which was produced using a 20% PCL solution at 15 kV. The physicochemical properties of the electrospun PCL scaffold are summarized in Table 1. The mean diameter of the fibers (2.08 ± 1.07 μm) fell within the range of the typical diameter of endogenous human collagen, which is 1–20 μm.^30^ The scaffold had adequate porosity—a property that promotes cell-surface interactions, nutrient and gas exchange, and cell proliferation^31^—with a mean pore size of 157 ± 173 μm^2^, which can sufficiently facilitate MSC infiltration. Despite its porous architecture, the scaffold maintained a high level of tensile strength at 9.38 ± 2.04 MPa.

**Figure 1.**
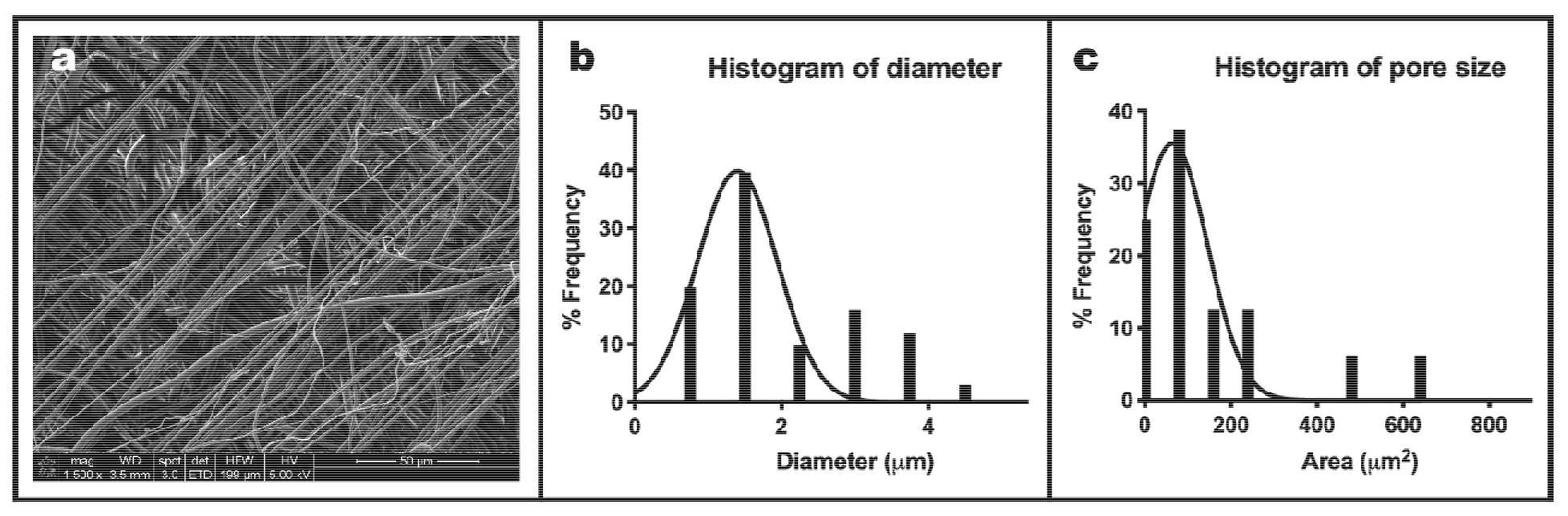
Alignment, fiber diameter, and pore size of electrospun PCL. (a) On scanning electron microscopy, the fibers of the electrospun scaffold showed good alignment (bar = 50 μm, 1 500x magnification). (b, c) Histograms showing the frequency distribution of the fiber diameters and scaffold pore sizes.

**Table 1.** Physicochemical properties of the electrospun PCL scaffold.

| Properties | PCL |
| --- | --- |
| Diameter ($\mu\text{m}$) | $2.08 \pm 1.07$ |
| Pore size ($\mu\text{m}^2$) | $157 \pm 173$ |
| Porosity (calculated, %) | $70.36 \pm 4.90$ |
| Porosity (intrusion, %) | $83.96 \pm 1.61$ |
| $T_g$ ($^{\circ}\text{C}$) | $-65.64 \pm 1.89$ |
| $T_m$ ($^{\circ}\text{C}$) | $55.73 \pm 0.77$ |
| Maximum stress (MPa) | $9.38 \pm 2.04$ |

Abbreviations: PCL, polycaprolactone; $T_g$ , glass transition temperature; $T_m$ , melting temperature.

### Electrospun PCL supported the attachment of MSCs

We have shown in a previous study that an electrospun PCL scaffold can support the attachment of human GFP-expressing MSCs, with the number of viable cells attached to the scaffold increasing as much as 7-fold after 1 week of incubation.^23^ In this study, fluorescence microscopy similarly revealed that the seeded Sprague-Dawley rat RFP-MSCs attached to the electrospun PCL scaffold (Figure 2).

**Figure 2.**
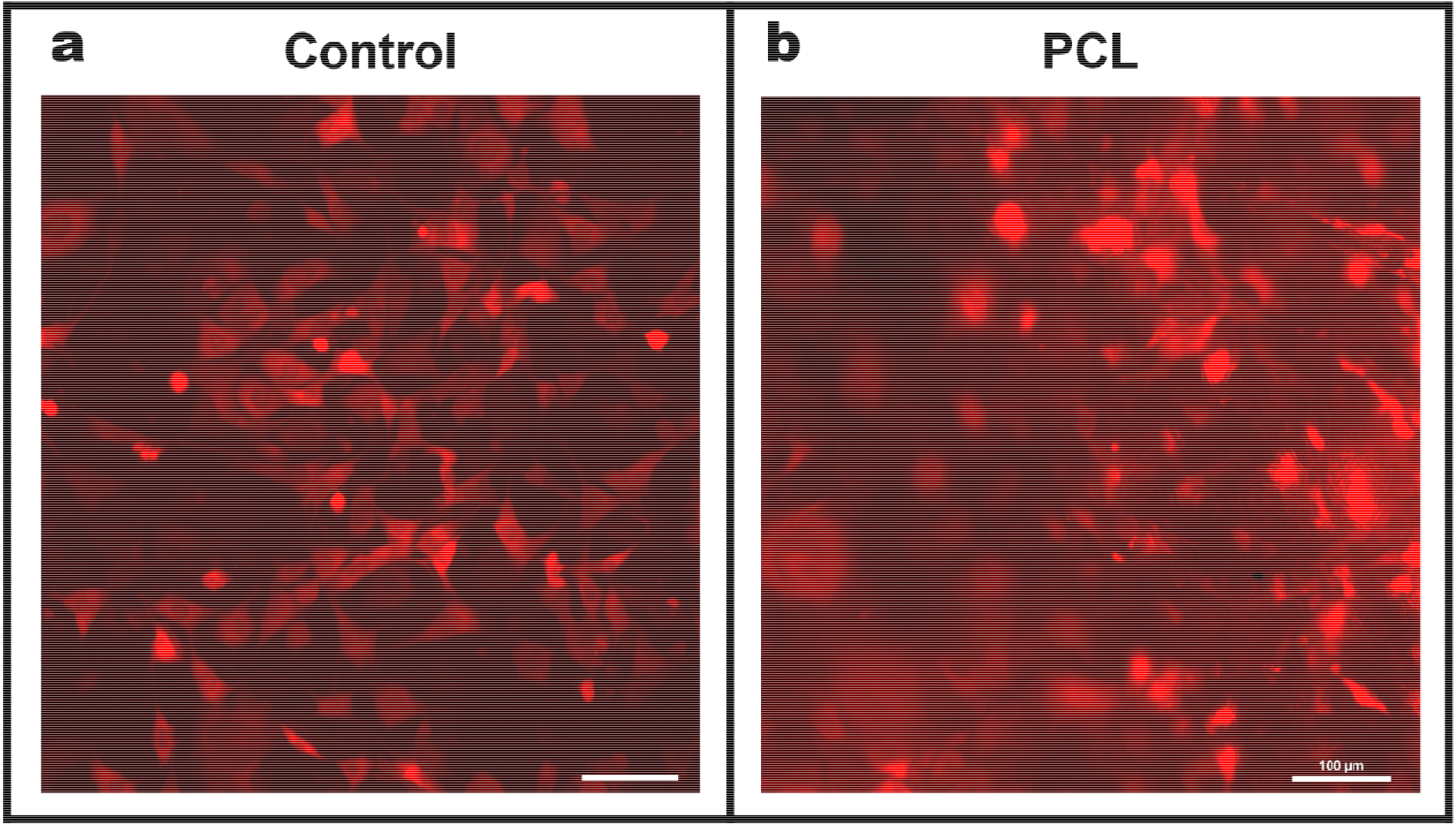
Attachment of Sprague-Dawley rat MSCs onto the electrospun PCL. (a) Control MSCs cultured on a polystyrene plate and (b) Sprague-Dawley rat RFP-MSCs loaded onto electrospun PCL scaffold after 12 hours of incubation and washing (bars = 100 μm, 10x magnification). Abbreviations: RFP-MSC, red fluorescent protein–mesenchymal stem cell; PCL, polycaprolactone.

### Successful CKD modeling in Sprague-Dawley rats

Representative images of the surgical procedures are shown in Figure 3. Impaired kidney function in the Sprague-Dawley rats was confirmed by increased serum creatinine and BUN. Four weeks after the second step of the two-stage surgery (i.e., right nephrectomy), the mean serum creatinine level of the CKD rats (N = 6, 0.83 ± 0.09 mg/dL) was 1.41 times higher than that of control rats (N = 3, 0.59 ± 0.13 mg/dL, *P* < 0.03), and the mean BUN level of the CKD rats (N = 6, 49.28 ± 6.99 mg/dL) was 1.54 times higher than that of the control rats (N = 3, 32.03 ± 2.67 mg/dL, *P* < 0.002).

**Figure 3.**
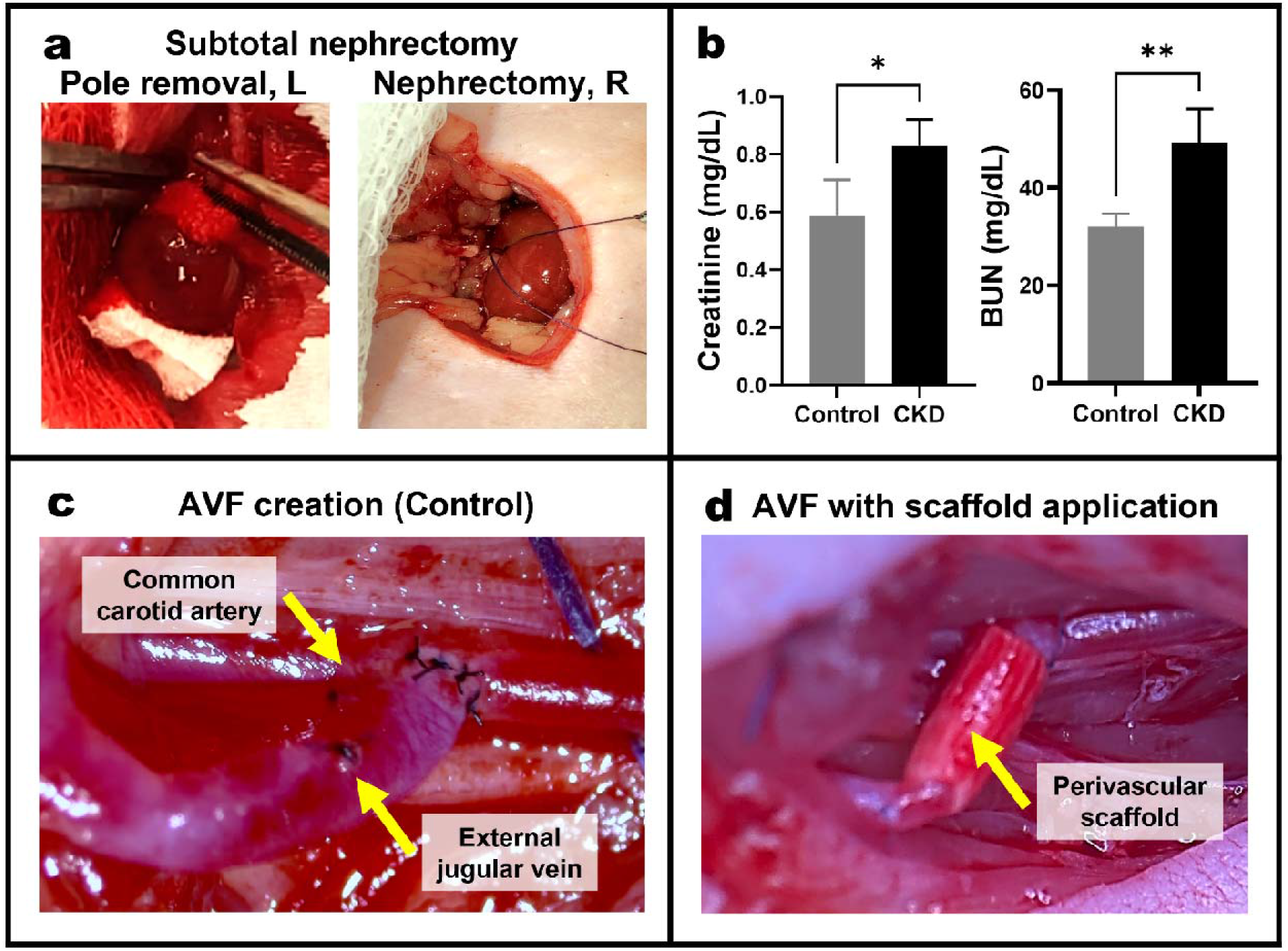
Surgical procedures and kidney function after 5/6^th^ nephrectomy. (a) Two-stage surgery to induce CKD in Sprague-Dawley rats. In step 1 (left image), the upper and lower poles of the left kidney are removed. In step 2 (right image), the right nephrectomy is performed 1 week later. (b) Significantly increased serum creatinine and BUN levels are observed 4 weeks following the two-stage surgery. (c/d) Surgical creation of AVFs via end-to-side anastomosis of the external jugular vein with the common carotid artery. Abbreviations: *, *P* < 0.03; **, *P* < 0.002; BUN, blood urea nitrogen; CKD, chronic kidney disease; L, left; R: right.

### AVFs of the CKD rats had greater NIH and inflammatory response than non-CKD rats

Figure 4 compares the AVFs in non-CKD and CKD rats after AVF creation. At 4 weeks, US measurements revealed that the mean wall thickness of the AVFs of the CKD rats (N = 6, 0.54 ± 0.14 mm) was 1.93 times higher than that of non-CKD rats (N = 6, 0.28 ± 0.11 mm, *P* < 0.002), while the mean luminal diameter of the AVFs of the CKD rats (N = 6, 0.74 ± 0.07 mm) was significantly lower than that of non-CKD rats (N = 6, 0.88 ± 0.09 mm, *P* < 0.03). Consequently, the wall-to-lumen ratio of the AVFs of the CKD rats (N = 6, 0.75 ± 0.22 mm) was 2.27 times higher than that of non-CKD rats (N = 6, 0.33 ± 0.14 mm, *P* < 0.002). In addition, on histomorphometric analysis, the AVFs of the CKD rats had a significantly higher neointima-to-lumen ratio, a measure of luminal stenosis, and a higher neointima-to-media ratio, a measure of wall thickening. The neointima-to-lumen ratio of the AVFs of the CKD rats (N = 3, 19.98 ± 5.84) was 31.22 times higher than that of non-CKD rats (N = 3, 0.64 ± 0.07, *P* < 0.002). The neointima-to-media ratio of the AVFs of the CKD rats (N = 3, 3.08 ± 0.05) was 6.84 times higher than that of non-CKD rats (N = 3, 0.45 ± 0.07, *P* < 0.0002). Moreover, when we measured inflammatory response via ^18^F-FDG PET imaging, the AVFs of the CKD rats had significantly higher ^18^F-FDG uptake as assessed by SUV_max_ (N = 6, 6.49 ± 1.31) compared to that of non-CKD rats (N = 4, 1.18 ± 0.51, *P* < 0.05). Therefore, in the succeeding experiments, CKD rats were used exclusively to determine the efficacy of the MSC-loaded scaffolds.

**Figure 4.**
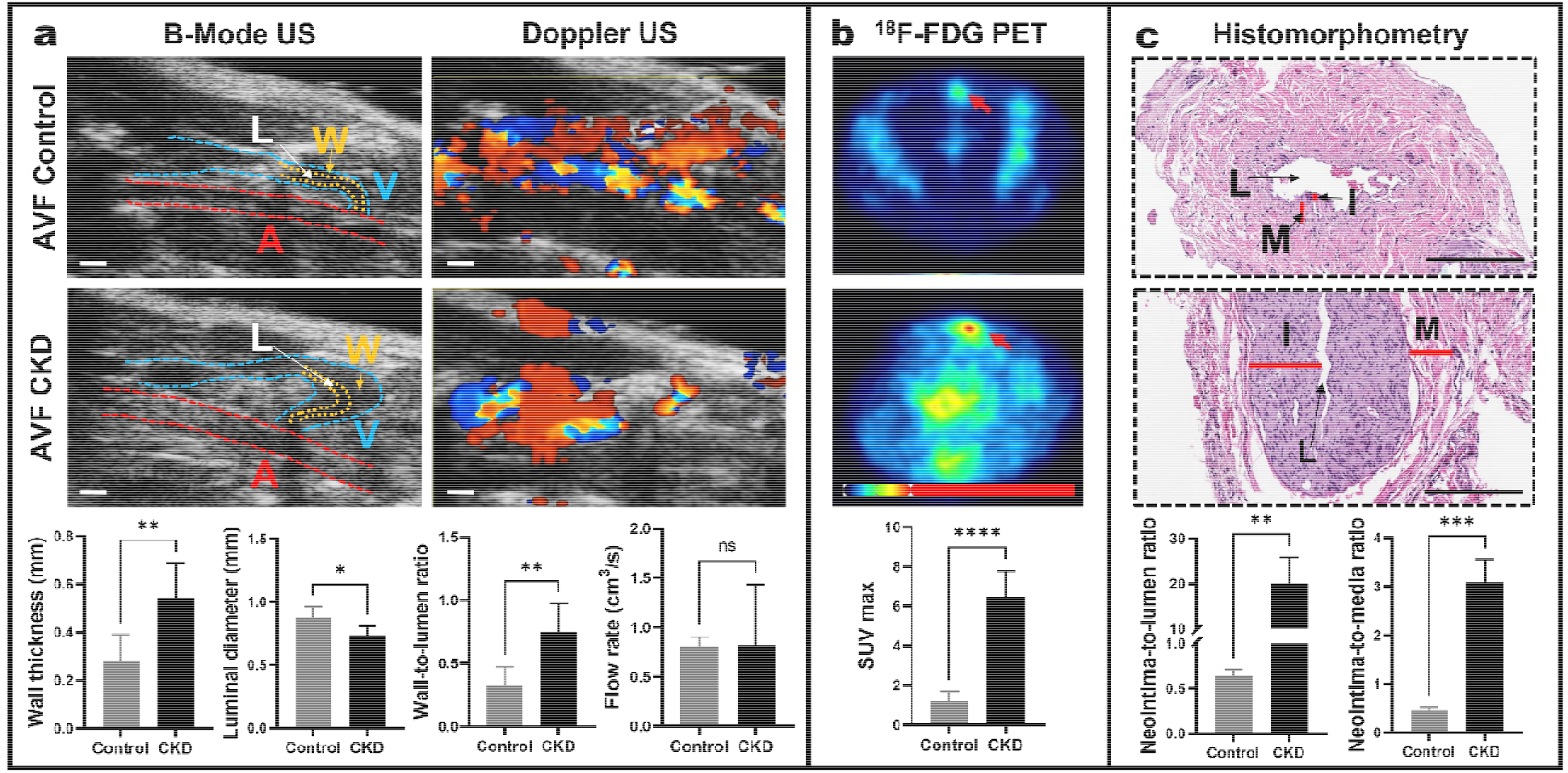
Comparison of non-CKD and CKD rats. (a) B-mode and color Doppler ultrasonography of the AVFs in non-CKD (i.e., control) and CKD rats. CKD rats had significantly higher mean wall thickness, lower luminal diameter, and higher wall-to-lumen ratio (A, artery; L, lumen; V, vein; W, wall; bars = 1 mm). (b) ^18^F-FDG-PET images demonstrate significantly higher SUV_max_ values at the outflow vein in CKD rats compared to non-CKD rats. (c) On histomorphometric analysis, neointima-to-lumen and neointima-to-media ratios were higher for CKD rats than for non-CKD rats (bars = 100 μm, 20x magnification; I, intima; L, lumen; M, media). Abbreviations: *, *P* < 0.03; **, *P* < 0.002; ***, *P* < 0.0002; ****, *P* < 0.0001; AVF, arteriovenous fistula; CKD, chronic kidney disease; ^18^F-FDG-PET, ^18^F-fluorodeoxyglucose positron emission tomography; ns, not significant; US, ultrasonography.

### PCL-alone and PCL+MSC wraps improved AVF lumen diameter, wall-to-lumen ratio, and flow rate

Figure 5 compares the B-mode and color Doppler ultrasonographic findings for AVFs in CKD Sprague-Dawley rats at 4 weeks following AVF creation and placement of no wrap (i.e., control rats, N = 12), PCL wrap alone (N = 12), and PCL+MSC wrap (N = 12). The addition of a PCL wrap alone resulted, when compared with control rats, in significantly increased luminal diameter (1.25 ± 0.11 mm versus 0.82 ± 0.10 mm, *P* < 0.0001), reduced wall-to-lumen ratio (0.40 ± 0.05 versus 0.69 ± 0.17, *P* < 0.0001), and increased flow rate (2.95 ± 0.80 versus 1.32 ± 0.99 cm^3^/s, *P* < 0.03). Likewise, compared to controls, the addition of PCL+MSC wraps resulted in significantly increased luminal diameter (1.74 ± 0.29 mm, *P* < 0.0001), reduced wall-to-lumen ratio (0.29 ± 0.06, *P* < 0.0001), and increased flow rate (5.74 ± 2.36 cm^3^/s, *P* < 0.0001). When comparing PCL alone and PCL+MSC wraps, PCL+MSC resulted in a greater improvement in luminal diameter (*P* < 0.0001), wall-to-lumen ratio (*P* < 0.03), and flow rate (*P* < 0.0002) compared to PCL alone, suggesting that the addition of MSCs promotes better luminal expansion and blood flow through the outflow vein.

**Figure 5.**
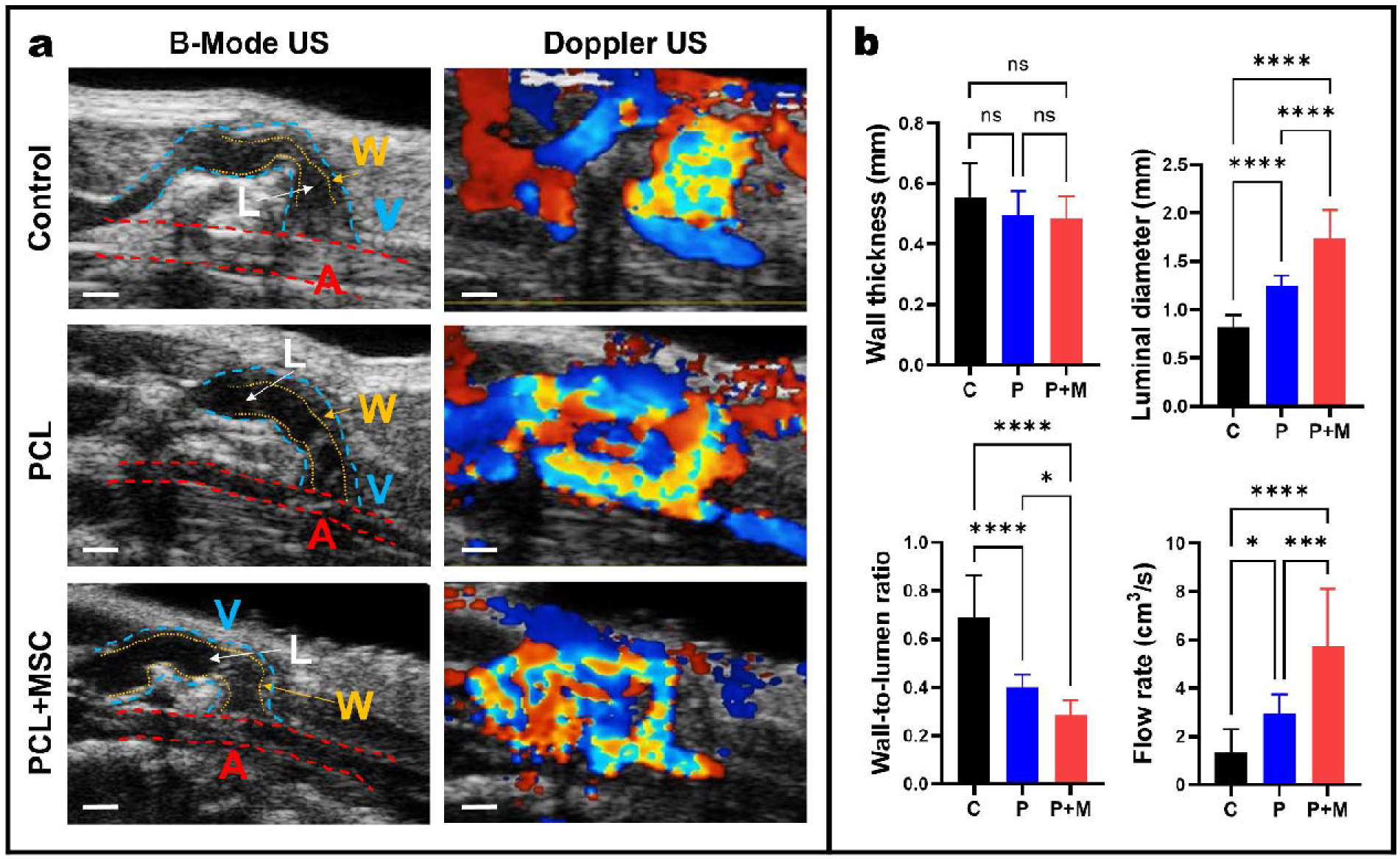
Evaluation of wall thickness, luminal diameter, and flow rate via B-mode and color Doppler ultrasonography. (a) Ultrasound images of the AVF in CKD rats with no wrap (i.e., control), PCL alone, and PCL+MSC perivascular wrap at 4 weeks (A, artery; L, lumen; V, vein; W, wall; bars = 1 mm). (b) Wall thickness, luminal diameter, wall-to-lumen ratio, and flow rate for the venous outflow in CKD rats with no wrap (i.e., control, C), PCL wrap alone (P), and PCL+MSC (P+M) wrap. While no significant difference was detected among the three groups with respect to wall thickness, significant increases in luminal diameter, decreases in wall-to-lumen ratio, and increases in flow rate were found when comparing C versus P, C versus P+M, and P versus P+M. Abbreviations: *, *P* < 0.03; **, *P* < 0.002; ***, *P* < 0.0002; ****, *P* < 0.0001; C, control; P/PCL, polycaprolactone; M/MSC, mesenchymal stem cell; ns, not significant; US, ultrasonography.

### PCL+MSC wrap reduced ^18^F-FDG-PET uptake at the outflow vein of AVFs in CKD rats

Figure 6 compares the PET findings for the AVFs of CKD rats with no wrap (i.e., control, N = 6), PCL-alone wrap (N = 6), and PCL+MSC wrap (N = 6) at 4 weeks after AVF creation. The perivascular application of PCL+MSC wrap resulted in significantly reduced ^18^F-FDG uptake compared to rats without no wrap (SUV_max_ 3.15 ± 1.96 versus 6.49 ± 1.31, *P* < 0.002). No significant difference was detected on ^18^F-FDG-PET imaging when comparing PCL-alone versus control CKD rats (SUV_max._ 7.07 ± 1.71 versus 6.49 ± 1.31, *P* = 0.8235).

**Figure 6.**
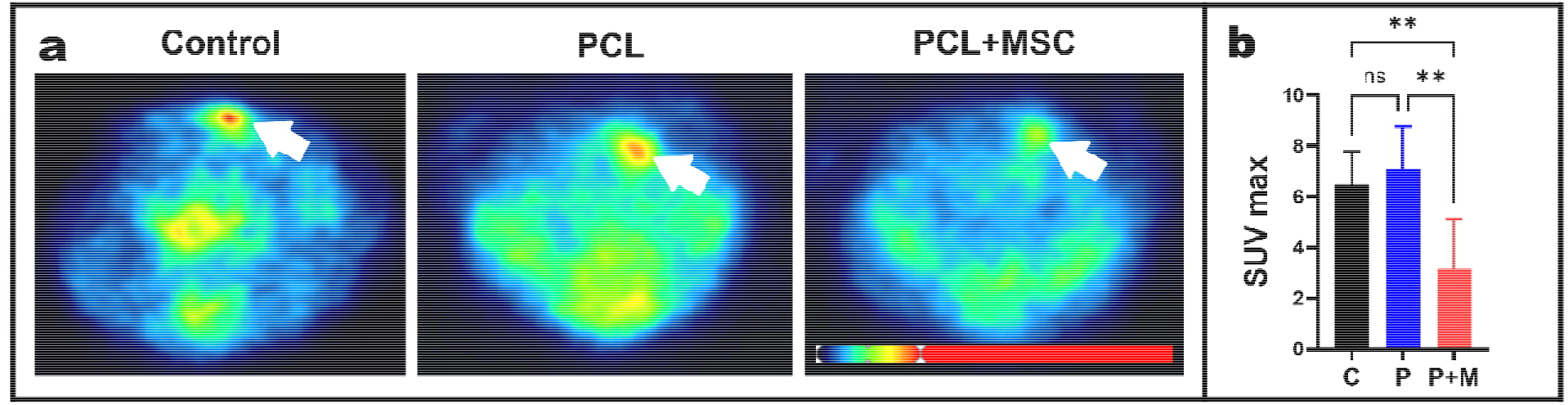
Axial ^18^F-FDG-PET images at the outflow vein of AVF in CKD rats. (a) PET images from rats with no wrap (i.e., control), PCL-alone wrap, and PCL+MSC wrap were obtained 4 weeks following surgery. The location of the fistula is marked by the white arrows. (b) SUV_max_ measurements show that PCL+MSC (P+M) resulted in a significant reduction in ^18^F-FDG uptake compared to control and PCL alone. Abbreviations: **, *P* < 0.002; C, control; P/PCL, polycaprolactone; M/MSC, mesenchymal stem cell; ns, not significant.

### PCL scaffold improved retention of RFP-MSCs

The presence of RFP-MSCs was investigated 2 and 4 weeks following AVF surgery in representative rats with no wrap (i.e., control), rats treated with PCL wrap, rats treated with a perivascular injection of RFP-MSCs, and rats treated with PCL+MSC wrap. An AVF treated with PCL+MSC qualitatively showed improved RFP-MSC retention compared to the AVF that received a perivascular injection of MSCs after 4 weeks, as seen on fluorescence microscopy (Figure 7).

**Figure 7.**
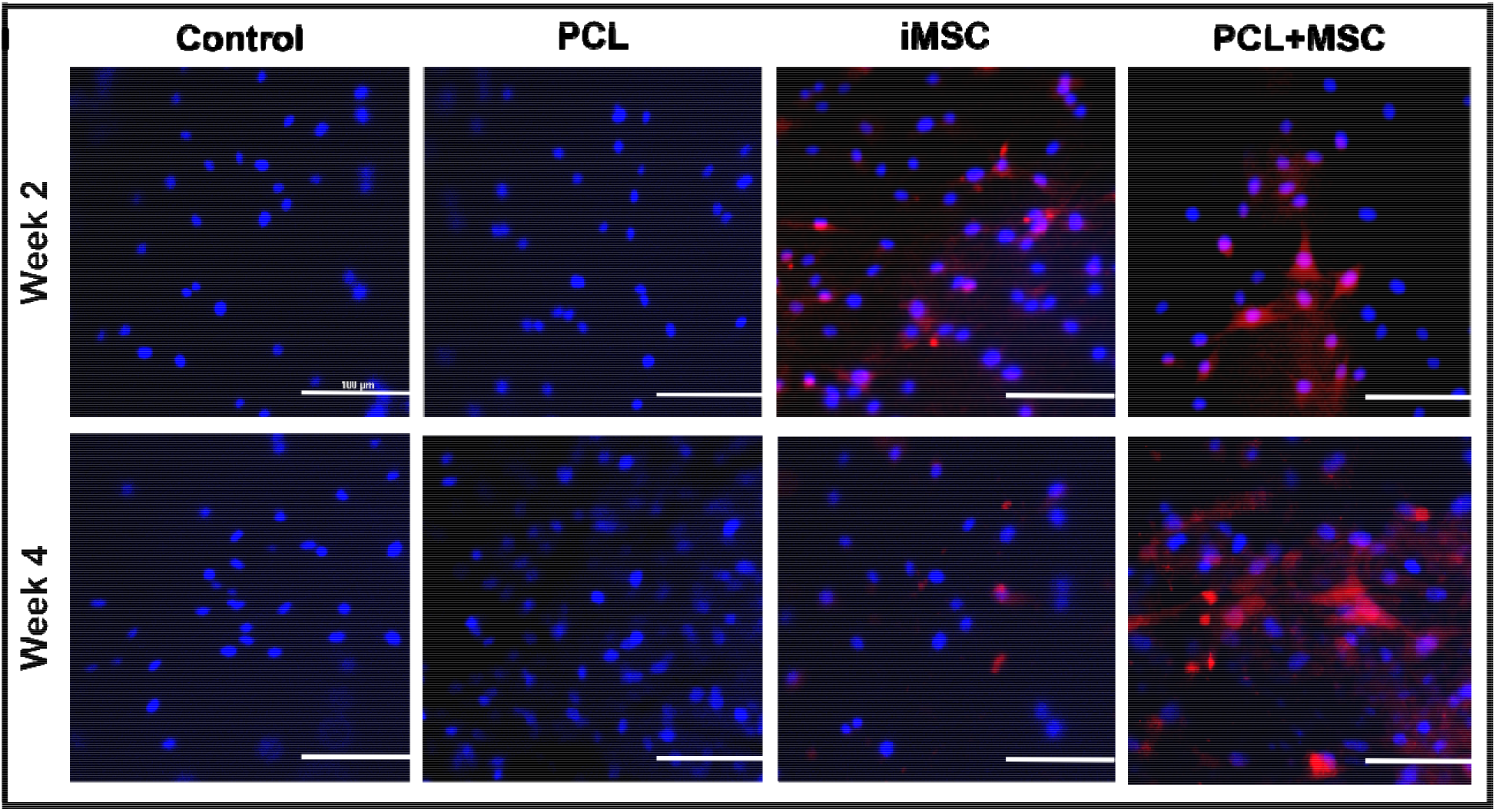
Fluorescence microscopy of the harvested AVFs of representative Sprague-Dawley CKD rats. Fluorescent microscopic images demonstrate the increased retention of RFP-MSCs at 2 and 4 weeks following surgery in the AVF that received PCL+MSC perivascular scaffold compared to perivascular injection of MSCs (iMSC) (bar = 100 μm). Abbreviations: PCL, polycaprolactone; MSC, mesenchymal stem cell.

### PCL-alone and PCL+MSC wraps reduced NIH at the outflow vein of AVF in CKD rats

Figure 8 compares the histological findings for the AVFs of CKD rats with no wrap (control, N = 3), PCL-alone wrap (N = 3), and PCL+MSC wrap (N = 3) at 4 weeks after AVF creation. The application of a PCL-alone wrap resulted, when compared to control rats, in a significantly reduced neointima-to-lumen ratio (0.26 ± 0.01 versus 19.98 ± 5.84, *P* < 0.0002) and neointima-to-media ratio (0.83 ± 0.05 versus 3.08 ± 0.47, *P* < 0.0002). Similarly, the application of a PCL+MSC wrap resulted in a significantly reduced neointima-to-lumen ratio (0.06 ± 0.01 versus 19.98 ± 5.84, *P* < 0.0002) and neointima-to-media ratio (0.17 ± 0.02 versus 3.08 ± 0.47, *P* < 0.0001) compared to controls. These findings confirm the beneficial effect of PCL-alone and PCL+MSC wraps on wall thickness and luminal diameter, which is concordant with the anatomic imaging findings obtained via ultrasonography. Although there was no significant difference detected between PCL-alone and PCL+MSC treatment in regard to neointima-to-lumen and neointima-to-media ratios, there was a trend towards further reduction in both these ratios with PCL+MSC treatment. When compared to the PCL-alone wrap, the PCL+MSC wrap reduced the mean neointima-to-lumen ratio and mean neointima-to-media ratio by 77% and 80%, respectively.

**Figure 8.**
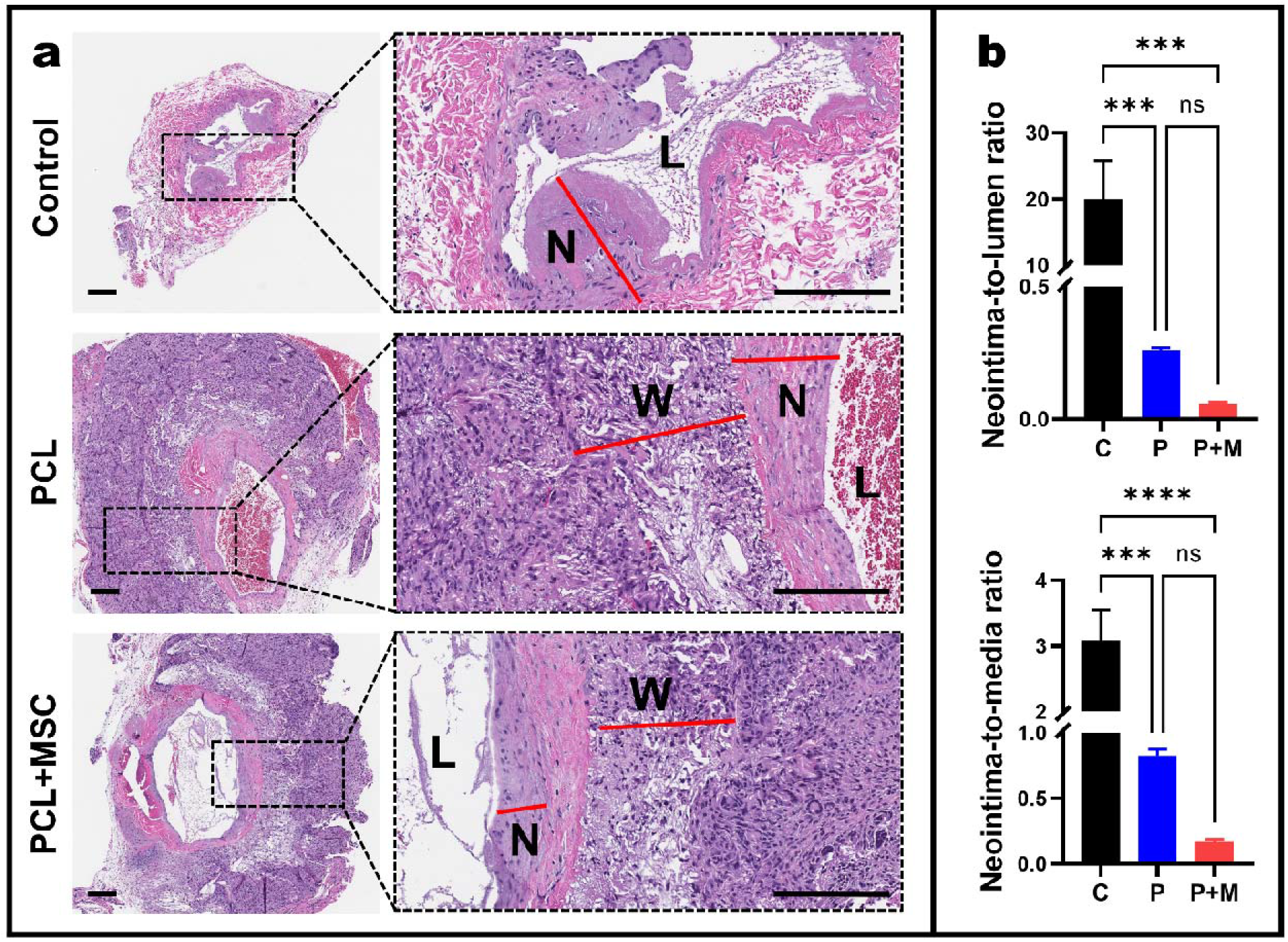
Histomorphometric analysis in Sprague-Dawley CKD rats 4 weeks following AVF surgery. (a) Histology of the outflow vein of AVF in rats with no wrap (i.e., control), PCL-alone wrap, and PCL+MSC wrap (bars = 100 μm, L: lumen; N: neointima; W: wrap). (b) Histomorphometry shows that both PCL and PCL+MSC perivascular wraps resulted in a significant reduction in both neointima-to-lumen and neointima-to-media ratios relative to control. Abbreviations: ***, *P* < 0.0002; ****, *P* < 0.0001; C, control; P/PCL, polycaprolactone; M/MSC, mesenchymal stem cell; ns, not significant.

The presence of cells positive for CD45, α-SMA, and vimentin was also investigated 4 weeks following AVF surgery in a representative rat with no wrap (i.e., control), rat treated with a perivascular injection of RFP-MSCs, and rat treated with PCL+MSC wrap. The AVFs treated with PCL and PCL+MSC revealed fewer cells that were positive for CD45, α-SMA, and vimentin (Figure 9, Table 2). These findings suggest the scaffold treatments can potentially reduce the neointimal infiltration of inflammatory cells, smooth muscle cells, and fibroblasts.

**Figure 9.**
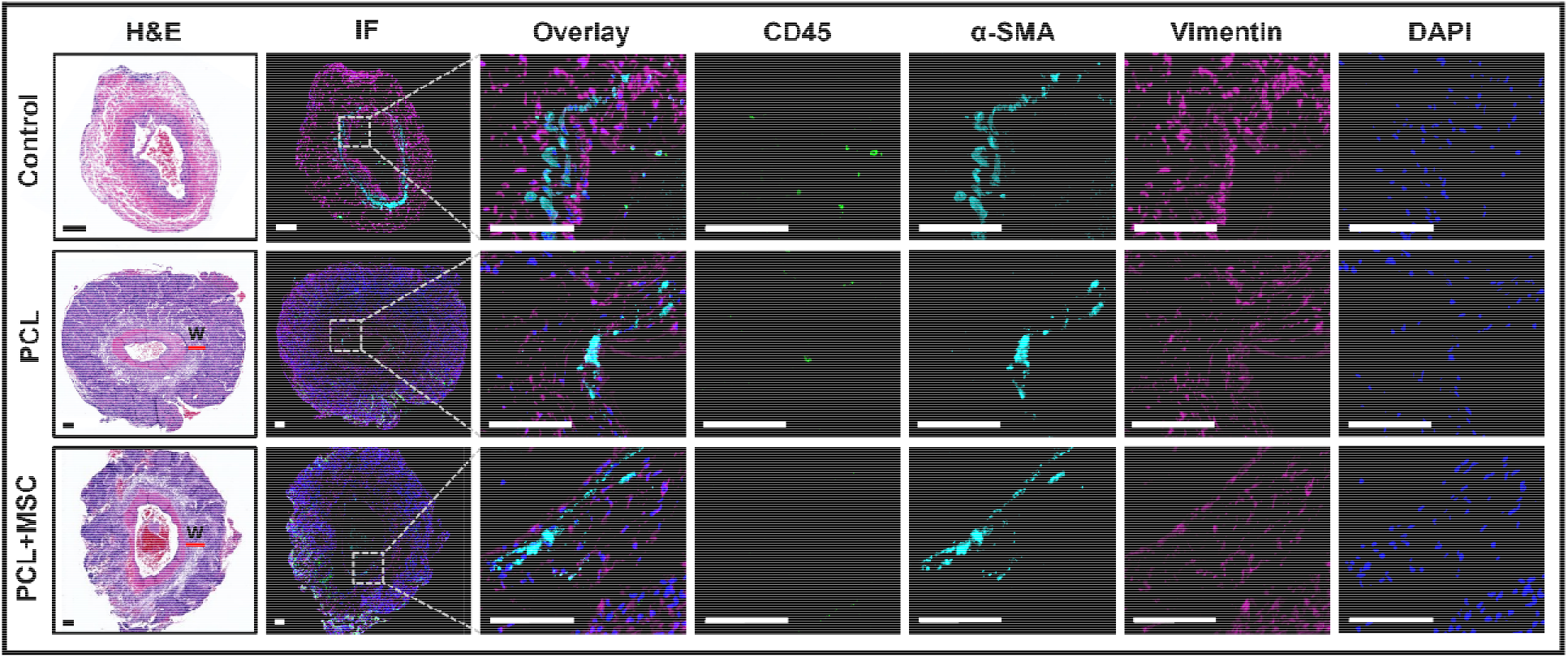
Immunofluorescence staining of the AVFs of representative Sprague-Dawley CKD rats. H&E with corresponding immunofluorescent microscopic images that demonstrate reduced CD45, α-SMA, and vimentin staining for both PCL and PCL+MSC perivascular wraps compared to no intervention (i.e., control) 4 weeks following surgery (bars = 100 μm, W: wrap). Abbreviations: α-SMA, alpha smooth muscle actin; CD, cluster of differentiation; DAPI, 4’,6-diamidino-2-phenylindole; H&E, hematoxylin, and eosin; IF, immunofluorescence; PCL, polycaprolactone; MSC, mesenchymal stem cell.

**Table 2.** Immunofluorescence staining analysis of the neointima.

| Marker | Control |  | PCL |  | PCL+MSC |  |
| --- | --- | --- | --- | --- | --- | --- |
|  | Cell count | % | Cell count | % | Cell count | % |
| CD45 | 37 | 24.03 | 1 | 0.83 | 1 | 0.57 |
| α-SMA | 97 | 62.99 | 30 | 24.79 | 31 | 17.61 |
| Vimentin | 60 | 38.96 | 47 | 38.84 | 44 | 25.00 |
| DAPI (Total) | 154 | 100.00 | 121 | 100.00 | 176 | 100.00 |

Abbreviations: α-SMA, alpha smooth muscle actin; CD, cluster of differentiation; DAPI, 4',6-diamidino-2-phenylindole; PCL, polycaprolactone; MSC, mesenchymal stem cell.

## DISCUSSION

In this study, we pioneered the use of a bioresorbable electrospun perivascular polymeric scaffold to provide mechanical strength to a maturing AVF and to support the attachment and proliferation of Sprague-Dawley rat MSCs. We optimized the polymer concentration and electrospinning voltage to produce scaffolds with aligned fibers, as previous work has shown that MSC-seeded scaffolds with highly aligned fibers have greater mechanical stability and yield higher degrees of cell proliferation compared to scaffolds with random orientation.^32,33^ Our scaffold also featured properties, such as fiber diameter, pore size, and porosity, that are comparable to those found in the endogenous human extracellular matrix.^30,31^ We anticipate that the constructed polymer is a highly suitable medium for the delivery of cell-based therapy for Sprague-Dawley rats, and potentially, for humans.

We found that the application of a PCL scaffold alone reduced the stenosis and thickening of the venous outflow wall of an AVF, as seen on US and histological analysis. These results indicate that mechanical support alone can improve the patency of a maturing AVF, which is concordant with prior research in humans.^18^ It should be noted that the optimal duration for perivascular wrap presence prior to complete dissolution is unknown. One of the major pathways that underlie NIH is hemodynamic stress. Specifically, disturbed or oscillatory wall shear stress leads to endothelial activation and, subsequently, endothelial proliferation. Moreover, mechanical loads that exceed the capacity of blood vessels eventually cause vascular wall injury and activation of synthetic smooth muscle cells and fibroblasts that proliferate and migrate centrally into the intima.^12^ Hence, the addition of a perivascular wrap, which could prevent the overexpansion of the vascular wall and reduce the flow turbulence within the vascular lumen, is likely to improve the patency of a maturing AVF.

We also found that incorporating MSCs onto the PCL scaffold yielded improvements in luminal diameter and flow rate and a significant reduction in ^18^F-FDG uptake, as seen on PET images. Moreover, the addition of MSCs to the PCL wraps resulted in a trend (nonsignificant) toward further reduction in the neointima-to-lumen and neointima-to-media ratios on H&E staining and a further reduction in the percentages of cells that were positive for CD45, α-SMA, and vimentin on immunofluorescence staining. Taken together, these findings suggest that mechanical interventions to promote AVF maturation and patency may be improved by the perivascular addition of MSCs. Given the rapid washout of perivascular injections of MSCs, the use of a perivascular wrap to implant MSCs, in turn, improves the efficacy of MSCs towards mitigating pathologic inflammation over time in the developing AVF.

Interestingly, we observed significant differences in the mean luminal diameter and mean flow rate of AVFs wrapped with a PCL scaffold alone versus an MSC-seeded PCL scaffold; such differences, however, were not reflected on histological specimens. This observation suggests there may be in vivo changes associated with eventual AVF maturation or, conversely, eventual AVF failure that occurs prior to visible changes in the composition and architecture of vascular tissue. Further investigation on the cellular and extracellular composition of the vascular wall is warranted to identify possible sources of variations between in vivo and histological observations. Although ^18^F-FDG-PET is associated with time and monetary costs, PET imaging offers a potential approach to estimating degrees of local inflammation within the AVFs in vivo. Hence, further studies are also warranted to identify the proper timing, frequency, and population for ^18^F-FDG-PET measurement in the setting of clinical AVF surveillance.

Overall, our results support the utilization of a perivascular scaffold to maximize the efficacy of MSCs to minimize pathologic NIH. Furthermore, our findings highlight the potential use of ^18^F-FDG-PET to identify early inflammatory changes following AVF formation, which could potentially be used as a non-invasive marker to assess patients at risk for pathologic inflammation and, therefore, at risk of AVF maturation failure. Although further studies are needed to evaluate the exact mechanisms by which the PCL-alone and PCL+MSC perivascular wraps mitigate NIH, our study provides a compelling basis for the development and clinical use of MSC-seeded perivascular scaffolds to improve AVF maturation and patency outcomes.

## AUTHOR CONTRIBUTIONS

Conceptualization, J.V.D.P., A.J.R.B., F.M.H., R.R.B., S.Y.H., M.P.M.; investigation, J.V.D.P., A.J.R.B., M.R.B., J.A.D., A.C., H.C.D.M., E.M.D.S.V., C.K., G.C., R.R.B.; data curation, J.V.D.P., A.J.R.B., M.R.B., J.A.D., A.C., H.C.D.M., E.M.D.S.V., C.K., G.C., R.R.B., M.P.M.; formal analysis, J.V.D.P., A.J.R.B., M.R.B., J.A.D., A.C., H.C.D.M., E.M.D.S.V., C.K., G.C., R.R.B., M.P.M.; writing—original draft, A.J.R.B., J.V.D.P.; writing—review and editing, A.J.R.B., J.V.D.P., M.R.B., J.A.D., A.C., H.C.D.M., E.M.D.S.V., C.K., G.C., F.M.H., R.A., N.F., R.R.B., J.C., S.Y.H., M.P.M.; visualization, J.V.D.P., A.J.R.B., M.R.B., J.A.D., A.C., H.C.D.M., E.M.D.S.V., C.K., R.R.B., M.P.M.; supervision, M.P.M.; project administration, M.P.M.; funding acquisition, F.M.H., S.Y.H., M.P.M.; All authors have read and agreed to the published version of the manuscript.

## ACKNOWLEDGMENTS

The authors would like to acknowledge Sunita C. Patterson in MD Anderson’s Research Medical Library for editing the manuscript, Dr. James Gu at the Electron Microscopy Core at Houston Methodist Research Institute for assisting with the conduct of scanning electron microscopy, Dunn Lab personnel (i.e., Amanda McWatters, Malea L. Williams, and Steve D. Parrish) for assisting with animal experiments, and Small Animal Imaging Facility personnel for assisting with animal imaging.

## DATA AVAILABILITY

The raw/processed data required to reproduce these findings cannot be shared at this time due to technical or time limitations. Data will be made available on request.

## SUPPLEMENTARY MATERIAL

**Supplementary Figure S1.**
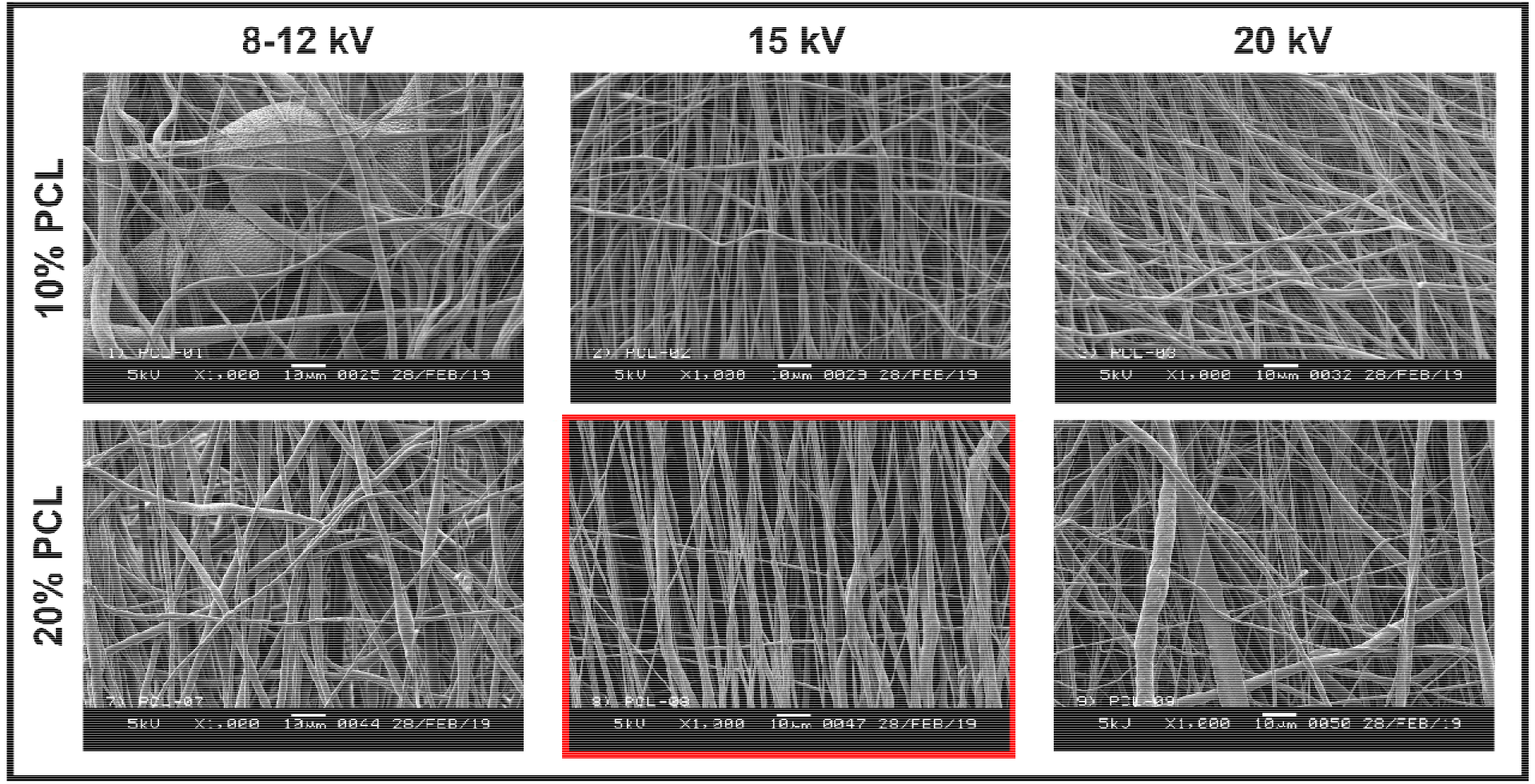
Optimization of polymer concentration and electrospinning voltage (bars = 10 μm, 1 000x magnification).

## REFERENCES

1. International Society of Nephrology. ISN Global Kidney Health Atlas. 2019. Published 2nd. Accessed September 15, 2022. https://www.theisn.org/initiatives/global-kidney-health-atlas/

2. United States Renal Data System. 2021 USRDS Annual Data Report: Epidemiology of kidney disease in the United States. National Institutes of Health, National Institute of Diabetes and Digestive and Kidney Diseases, Bethesda, MD, 2021 2021. Accessed September 15, 2022. https://adr.usrds.org/2021

3. Lok CE, Huber TS, Lee T, et al. KDOQI Clinical Practice Guideline for Vascular Access: 2019 Update. Am J Kidney Dis. 2020;75(4 Suppl 2):S1–S164.

4. Dember LM, Beck GJ, Allon M, et al. Effect of clopidogrel on early failure of arteriovenous fistulas for hemodialysis: a randomized controlled trial. JAMA. 2008;299(18):2164–2171.

5. Wasse H, Huang R, Naqvi N, Smith E, Wang D, Husain A. Inflammation, oxidation and venous neointimal hyperplasia precede vascular injury from AVF creation in CKD patients. J Vasc Access. 2012;13(2):168–174.

6. Simone S, Loverre A, Cariello M, et al. Arteriovenous fistula stenosis in hemodialysis patients is characterized by an increased adventitial fibrosis. J Nephrol. 2014;27(5):555–562.

7. Dixon BS, Beck GJ, Vazquez MA, et al. Effect of dipyridamole plus aspirin on hemodialysis graft patency. N Engl J Med. 2009;360(21):2191–2201.

8. Hammes M. Hemodynamic and biologic determinates of arteriovenous fistula outcomes in renal failure patients. Biomed Res Int. 2015;2015:171674.

9. Allon M, Robbin ML. Increasing arteriovenous fistulas in hemodialysis patients: problems and solutions. Kidney Int. 2002;62(4):1109–1124.

10. Lee T, Ullah A, Allon M, et al. Decreased cumulative access survival in arteriovenous fistulas requiring interventions to promote maturation. Clin J Am Soc Nephrol. 2011;6(3):575–581.

11. Lee T, Misra S. New Insights into Dialysis Vascular Access: Molecular Targets in Arteriovenous Fistula and Arteriovenous Graft Failure and Their Potential to Improve Vascular Access Outcomes. Clin J Am Soc Nephrol. 2016;11(8):1504–1512.

12. Brahmbhatt A, Remuzzi A, Franzoni M, Misra S. The molecular mechanisms of hemodialysis vascular access failure. Kidney Int. 2016;89(2):303–316.

13. Mohamed I, Kamarizan MFA, Da Silva A. Medical adjuvant treatment to increase patency of arteriovenous fistulae and grafts. Cochrane Database Syst Rev. 2021;7(7):Cd002786.

14. Barcena AJR, Perez JVD, Liu O, et al. Localized Perivascular Therapeutic Approaches to Inhibit Venous Neointimal Hyperplasia in Arteriovenous Fistula Access for Hemodialysis Use. Biomolecules. 2022;12(10):1367.

15. Prockop DJ, Olson SD. Clinical trials with adult stem/progenitor cells for tissue repair: let’s not overlook some essential precautions. Blood. 2007;109(8):3147–3151.

16. Han Y, Yang J, Fang J, et al. The secretion profile of mesenchymal stem cells and potential applications in treating human diseases. Signal Transduct Target Ther. 2022;7(1):92.

17. Yang B, Brahmbhatt A, Nieves Torres E, et al. Tracking and Therapeutic Value of Human Adipose Tissue-derived Mesenchymal Stem Cell Transplantation in Reducing Venous Neointimal Hyperplasia Associated with Arteriovenous Fistula. Radiology. 2016;279(2):513–522.

18. Chemla E, Velazquez CC, D’Abate F, Ramachandran V, Maytham G. Arteriovenous fistula construction with the VasQ™ external support device: a pilot study. J Vasc Access. 2016;17(3):243–248.

19. San Valentin EMD, Barcena AJR, Klusman C, Martin B, Melancon MP. Nano-embedded medical devices and delivery systems in interventional radiology. Wiley Interdiscip Rev Nanomed Nanobiotechnol. 2022:e1841.

20. Clair D, Moritz M, Burgess J, et al. Arteriovenous Fistula Outcomes After Local Vascular Delivery of a Sirolimus Formulation. J Vasc Surg. 2019;70(2):e37–e38.

21. Paulson WD, Kipshidze N, Kipiani K, et al. Safety and efficacy of local periadventitial delivery of sirolimus for improving hemodialysis graft patency: first human experience with a sirolimus-eluting collagen membrane (Coll-R). Nephrol Dial Transplant. 2012;27(3):1219–1224.

22. Siddiqui N, Asawa S, Birru B, Baadhe R, Rao S. PCL-Based Composite Scaffold Matrices for Tissue Engineering Applications. Mol Biotechnol. 2018;60(7):506–532.

23. Perez JVD, Singhana B, Damasco J, et al. Radiopaque scaffolds based on electrospun iodixanol/polycaprolactone fibrous composites. Materialia. 2020;14.

24. Robbin ML, Greene T, Allon M, et al. Prediction of Arteriovenous Fistula Clinical Maturation from Postoperative Ultrasound Measurements: Findings from the Hemodialysis Fistula Maturation Study. J Am Soc Nephrol. 2018;29(11):2735–2744.

25. Roberts JT, Villanueva-Meyer J, Bezold S, Krider SO, Nguyen QD. F-18-Fluorodeoxyglucose Positron Emission Tomography/CT Effectively Identifying Source of Infection in a Patient With Multiple Dialysis Arteriovenous Fistula Access Points. Cureus. 2020;12(6):e8516.

26. Zhu X, Cui W, Li X, Jin Y. Electrospun Fibrous Mats with High Porosity as Potential Scaffolds for Skin Tissue Engineering. Biomacromolecules. 2008;9(7):1795–1801.

27. Soliman S, Pagliari S, Rinaldi A, et al. Multiscale three-dimensional scaffolds for soft tissue engineering via multimodal electrospinning. Acta Biomater. 2010;6(4):1227–1237.

28. Wang X, Chaudhry MA, Nie Y, Xie Z, Shapiro JI, Liu J. A Mouse 5/6th Nephrectomy Model That Induces Experimental Uremic Cardiomyopathy. J Vis Exp. 2017(129):55825.

29. Wong CY, de Vries MR, Wang Y, et al. A Novel Murine Model of Arteriovenous Fistula Failure: The Surgical Procedure in Detail. J Vis Exp. 2016(108):e53294.

30. Ushiki T. [The three-dimensional ultrastructure of the collagen fibers, reticular fibers and elastic fibers: a review]. Kaibogaku Zasshi. 1992;67(3):186–199.

31. Green JJ, Elisseeff JH. Mimicking biological functionality with polymers for biomedical applications. Nature. 2016;540(7633):386–394.

32. Teh TK, Toh SL, Goh JC. Aligned hybrid silk scaffold for enhanced differentiation of mesenchymal stem cells into ligament fibroblasts. Tissue Eng Part C Methods. 2011;17(6):687–703.

33. Wang Z-y, Teoh SH, Johana NB, et al. Enhancing mesenchymal stem cell response using uniaxially stretched poly(ε-caprolactone) film micropatterns for vascular tissue engineering application. J Mater Chem B. 2014;2(35):5898–5909.

